# Structural basis underlying specific biochemical activities of non-muscle tropomyosin isoforms

**DOI:** 10.1101/2022.05.12.491677

**Authors:** Muniyandi Selvaraj, Shrikant Kokate, Gabriella Reggiano, Konstantin Kogan, Tommi Kotila, Elena Kremneva, Frank DiMaio, Pekka Lappalainen, Juha T. Huiskonen

**Affiliations:** Institute of Biotechnology, Helsinki Institute of Life Science HiLIFE, P.O Box 56, 00014 University of Helsinki, Helsinki, Finland; Department of Biochemistry, University of Washington, Seattle, WA 98195, USA

**Keywords:** actin, non-muscle tropomyosin, cryo-EM, cofilin, cytoskeleton

## Abstract

The actin cytoskeleton is critical for cell migration, morphogenesis, endocytosis, organelle dynamics, and cytokinesis. To support diverse cellular processes, actin filaments form a variety of structures with specific architectures and dynamic properties. Key proteins specifying actin filaments are tropomyosins. Non-muscle cells express several functionally non-redundant tropomyosin isoforms, which differentially control the interactions of other proteins, including myosins and ADF/cofilin, with actin filaments. However, the underlying molecular mechanisms have remained elusive. By determining the cryogenic electron microscopy structures of actin filaments decorated by two functionally distinct non-muscle tropomyosin isoforms, Tpm1.6 and Tpm3.2, we reveal that actin filament conformation remains unaffected upon binding. However, Tpm1.6 and Tpm3.2 follow different paths along the major groove of the actin filament, providing an explanation for their incapability to co-polymerize on actin filaments. The structures and biochemical work also elucidate the molecular basis underlying specific roles of Tpm1.6 and Tpm3.2 in myosin II activation and protecting actin filaments from ADF/cofilin-catalysed severing.

## INTRODUCTION

The actin cytoskeleton is critical for various cellular processes such as morphogenesis, motility, endocytosis, mechanosensing, and cytokinesis. To fulfill the specific needs of these diverse cellular processes, actin filaments assemble into a variety of structures with different architectures and dynamic properties and produce force through coordinated filament polymerization, as well as by serving as tracks for myosin motor proteins. Actin filaments of the different cytoskeletal structures associate with specific sets of actin-binding proteins, which give rise to distinct properties of actin filaments (Blanchoin et al., 2014; Boiero Sanders et al., 2020). In plants, the functional variety of actin filament structures has been proposed to derive from the presence of a large number of closely related actin isoforms (Gunning et al., 2015a). In contrast, fungi and metazoan cells express only one or few actin isoforms, and thus the functionally different actin filament structures in these organisms rely on proteins interacting with actin. These include tropomyosins (Tpms), which are elongated α-helical dimers that form head-to-tail oligomers along actin filaments. Tpms control the interactions of other proteins with actin filaments and regulate the stability of the actin filaments (Gunning et al., 2015b).

The functions of Tpms have been studied extensively in striated muscles. Tpms, together with a heteromeric protein complex consisting of troponin T, troponin I, and troponin C, control the association of myosin motor II domains of thick filaments with thin actin filaments in a Ca^2+^-dependent manner (Geeves, 2012). Recent cryogenic electron microscopy (cryo-EM) structures shed light on the mechanisms by which muscle Tpms (Tpm1.1 homodimer or Tmp1.1/Tpm2.2. heterodimers) interact with muscle actin filaments, and how the Tpm–troponin complex controls association of myosin II motor domains with actin filaments in muscle sarcomeres (Behrmann et al., 2012; Doran et al., 2020; Risi et al., 2021; Von der Ecken et al., 2015; Yamada et al., 2020). In an inactive Ca^2+^-free state (also referred to as blocked or ‘B-state’), the troponin complex stabilizes Tpm on an actin filament in a position that hinders myosin motor domain interaction. In the presence of Ca^2+^, the troponin complex undergoes a conformational change, which shifts the Tpm oligomer within the major groove of the actin filament by ∼10 Å to the ‘C-state’, where the myosin-binding site is partially uncovered (Yamada et al., 2020). Myosin-binding to the actin filament further moves Tpm to an otherwise energetically unfavorable ‘M-state’ position on an actin filament, resulting in actin-mediated activation of the myosin ATPase (Behrmann et al., 2012; Doran et al., 2020; Risi et al., 2021). In the absence of troponin complex and other proteins (apo or ‘A-state’), Tpm binds to an intermediate position between the B-and C-states that overlaps with the myosin-binding site (Von der Ecken et al., 2015).

Tropomyosins play important roles also in non-muscle cells where they can activate myosins similarly to striated muscles, as well as inhibit actin-interactions of other proteins such as ADF/ cofilin and fimbrin (Christensen et al., 2017; Coulton et al., 2010; Ono and Ono, 2002). Consequently, in animal non-muscle cells, Tpms are critical for the maintenance and function of actomyosin structures (Kumari et al., 2020; Tojkander et al., 2011). A large number (>40) of different Tpm isoforms can be generated through alternative splicing from four *TPM* genes in mammals, and several different Tpm isoforms are typically co-expressed in non-muscle cells (Meiring et al., 2019b). Depending on their length, these proteins can be classified into high- and low-molecular-weight Tpms. Different non-muscle Tpm isoforms typically display at least partially non-overlapping localization patterns in cells and are functionally non-redundant with each other (Bach et al., 2009; Bryce et al., 2003; Cagigas et al., 2021; Gallant et al., 2011; Johnson et al., 2014; Meiring et al., 2019a; Tojkander et al., 2011). Moreover, biochemical studies on a subset of the most abundant non-muscle Tpm isoforms revealed that they differentially affect ADF/cofilin-mediated actin filament disassembly and activation of non-muscle myosin II. For example, Tpm1.6 and Tpm1.7 inhibit ADF/cofilin-catalysed actin filament severing but do not activate myosin II. On the other hand, Tpm3.1, Tpm3.2, and Tpm4.2 do not efficiently protect actin filaments from ADF/cofilin but increase the non-muscle myosin II steady-state ATPase rate (Gateva et al., 2017; Jansen and Goode, 2019; Pathan-Chhatbar et al., 2018). Despite the importance of different Tpm isoforms in specifying functionally distinct actin filaments in non-muscle cells, the underlying molecular mechanisms remain elusive.

We hypothesized that the functional differences of the non-muscle tropomyosins may arise from their different effects on the conformation of the actin filament they interact with, or because that their interactions with the actin filament, and thus their positions relative to it, are different. The distinct functions could also be, at least partially, due to differences in the amino acid sequences of Tpms that could modulate the binding of other proteins on Tpm-decorated actin filaments. To address these different hypotheses, we focused here on two functionally distinct members of the family, Tpm1.6 and Tpm3.2. Of these two, Tpm1.6 exhibits a very slow off-rate from actin filaments, protects filaments from ADF/cofilin-catalysed severing but does not activate non-muscle myosin II. Tpm3.2, on the other hand, displays rapid dynamics on actin filaments and accelerates the steady-state ATPase rate of non-muscle myosin II in the presence of actin filaments, but does not efficiently protect actin filaments from ADF/cofilin (Gateva et al., 2017). By determining the cryogenic electron microscopy (cryo-EM) structures of β/γ-actin filaments decorated either by Tpm1.6 or by Tpm3.2, we revealed that the conformation of the actin filament remains unaffected. However, the two non-muscle Tpm isoforms follow slightly different paths along the major groove of the actin filament. Molecular modeling provides a plausible explanation for the incapability of these two Tpm isoforms to co-polymerize with each other on actin filaments and for their different effects on myosin II and ADF/cofilins. This is further corroborated by our biochemical assays. Taken together, our results shed light on the specific biochemical activities of non-muscle tropomyosins.

## RESULTS

### Structures of β/γ-actin filaments decorated with non-muscle tropomyosins

To determine the molecular architecture of functionally distinct non-muscle actin– tropomyosin complexes, we expressed and purified Tpm1.6 and Tpm3.2 isoforms. Tpm1.6 is a high-molecular-weight Tpm (284 residues) generated from the *TPM1* gene, whereas Tpm3.2 is a low-molecular-weight Tpm (248 residues) generated from the *TPM3* gene. The two isoforms display high amino acid sequence conservation in their central regions but differ especially in the regions adjacent to their ends (Fig. S1). We performed cryo-EM imaging on reconstituted complexes of both Tpm1.6 and 3.2 with β/γ-actin filaments. Our initial attempts to image the complexes revealed only bare actin, although either Tpm1.6 or Tpm3.2 had been added in excess. However, replacing NaCl in the sample buffer with sodium acetate (see Methods), as previously done in studies on muscle actin:Tpm:myosin complexes (Doran et al., 2020), resulted in a good decoration of actin filaments by both non-muscle Tpms.

Using a single-particle-based helical reconstruction approach (He and Scheres, 2017), we reconstructed the 3D structure of the actin:Tpm1.6 complex at an average resolution of 3.9 Å (Fig 1A,B, Fig. S2, Table S1). The filament measures approximately 90 Å in diameter (Fig. 1A). Actin was well-resolved, allowing atomic model building. The Tpm1.6 density was ordered sufficiently to separate the coiled-coils formed by the two α-helices in the Tpm1.6 dimer (Fig. 1A,B). However, the symmetry mismatch in the actin:Tpm1.6 assemblies, owing to the binding of one Tpm molecule per several actin monomers, hampered resolving the side chains of Tpm1.6 amino acid residues in the averaged density (Von der Ecken et al., 2015). Furthermore, the backbones of Tpm1.6 α-helices in the coiled-coil were unresolved, suggesting that the turn of the α-helix is not in phase with the symmetry of the actin filament, resulting in incoherent averaging of the Tpm1.6 backbone. Notably, the binding of Tpm1.6 left the structure of the actin filament unaltered; the refined helical symmetry parameters of actin:Tpm1.6 (rise 27.8 Å; turn: –166.5°) were nearly identical to those of bare chicken muscle actin (27.5 Å, –166.6°) reported earlier (Chou and Pollard, 2019).

**Figure 1.**
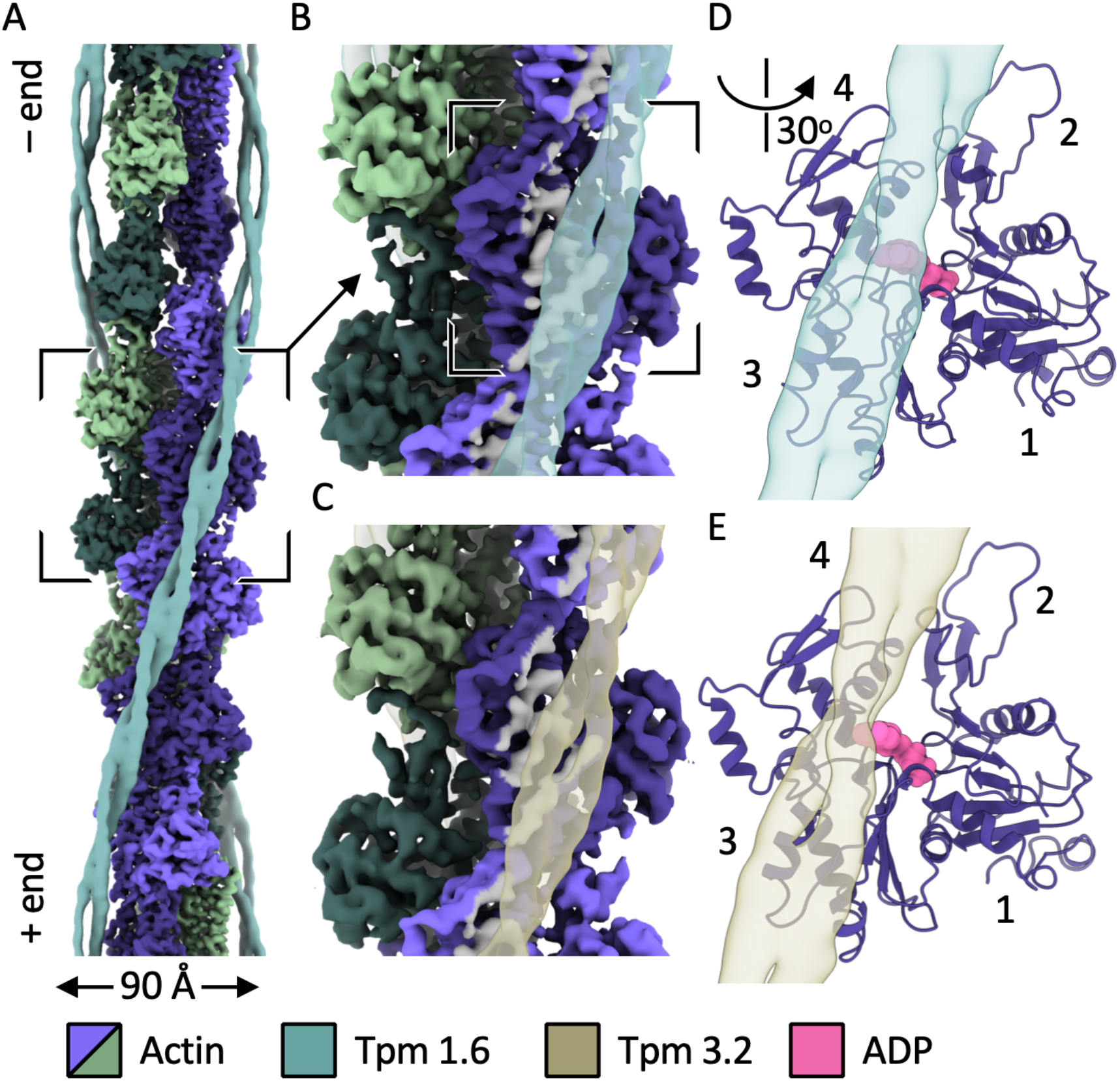
Cryo-EM structures of actin/non-muscle tropomyosin complexes. (**A**) Cryo-EM density of actin:Tpm1.6 complex is shown. Actin plus (barbed) and minus (pointed) ends are labelled. The density has been segmented and actin subunits are colored individually. The Tpm1.6 density has been filtered to 7-Å resolution. (**B**) A close-up of the area indicated in *A* is shown. Tmp1.6 density is shown as a transparent surface. Those parts of the actin surface that reside at 10 Å distance or closer to the Tpm1.6 have been colored in gray. (**C**) The same view as in *b* is shown for actin:Tpm3.2 complex. (**D**) A close-up of the area highlighted in *B* is shown rotated as indicated. The actin model is shown as a ribbon, the ADP model is shown as a surface, and the Tpm1.6 density as a transparent surface. The different sub-domains of actin are labelled 1–4. (**E**) The same view as in *d* is shown for actin Tpm3.2 complex. Tpm3.2 is shifted slightly in its position relative to the Tpm1.6 density in *D*.

To compare the binding of Tpm3.2 and Tpm1.6 to actin filaments, we also determined the structure of actin:Tpm3.2 complex to 4.6-Å resolution, using the same method as above (Fig. 1C; Fig. S3; Supplementary Table S1). A comparison of the Tpm densities relative to actin in these two complexes showed that the Tpm1.6 and Tpm3.2 dimers bind on different tracks on the surface of the actin filament (Fig. 1B,C). In both complexes, the Tpm binds to the major groove region of actin involving actin subdomains 1, 3, and 4, but excluding residues of actin subdomain 2 (Fig 1D,E). However, the position of Tpm3.2 is shifted slightly (∼5 Å) toward subdomains 3 and 4 when compared to the position of Tpm1.6. The root-mean-square-deviation (RMSD) was 1.02 Å for Tpm1.6-decorated actin (between 364 Cα-atom positions) and 1.08 Å for Tpm3.2-decorated actin (between 365 Cα-atom positions) when compared to bare actin (PDB:6DJO) (Chou and Pollard, 2019), indicating that all three actin structures have a highly similar conformation. The helical parameters of Tpm3.2-decorated actin (rise 27.8 Å, turn – 166.4°) were also nearly identical to those of bare actin. Together, these structures reveal that the two functionally distinct non-muscle Tpm isoforms leave the conformation of the actin filament unaffected but follow slightly different paths along the major groove of the actin filament.

### Two non-muscle tropomyosin isoforms bind actin via analogous salt bridges

In the absence of cryo-EM density for tropomyosin side chains, we turned to atomistic modeling using Rosetta to study the detailed interactions of Tpm1.6 and Tpm3.2 with actin filaments. We modeled the two Tpms as coiled-coils by combining AlphaFold with Rosetta-based energy minimization using the cryo-EM density maps. The interfaces generated by Alphafold were verified by Rosetta threading (see Methods). The longer Tpm1.6 and shorter Tpm3.2 models span the distance of about seven and six actin monomers, respectively (Fig. 2).

**Figure 2.**
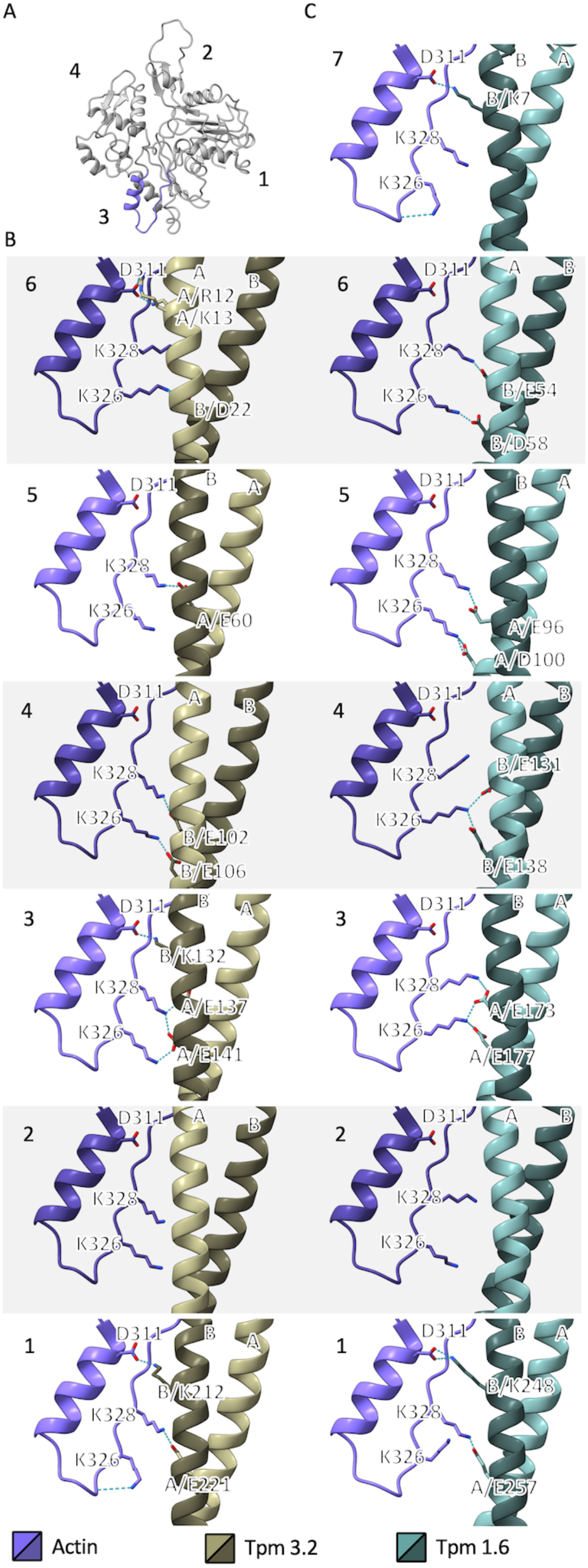
Models of the actin/non-muscle tropomyosin binding sites. (**A**) An atomic model of actin from actin:Tpm1.6 complex is shown as a ribbon. The different sub-domains of actin are labeled 1–4. The helix-and-loop motif shown in close-up in *B* and *C* is in blue. (**B**) Models of the binding sites between six actins and a single Tpm3.2 coiled-coil are shown. The two alpha-helices of the coiled-coil are labelled A and B. Sidechains are shown for actin residues D311, K326, and K328. Sidechains are also shown for those Tpm3.2 residues that are at hydrogen bonding distance. Hydrogen bonds predicted by the model are shown with dashed lines (cyan). (**C**) The same rendering as in *B* is shown for seven actins and Tpm1.6. Please not that one of the interactions is absent in Tpm3.2 due to its shorter length.

To study the charge complementarity between actin and Tpms, we calculated their electrostatic surfaces together with that of actin. A positively charged region on actin resides in the proximity of the Tpm density (Fig. S4). It corresponds to residues Lys326 and Lys328, which have been predicted to bind tropomyosins via salt bridges, alongside negatively charged Asp311 (Lehman et al., 2019; Pavadai et al., 2020) (see below). Most of the actin-facing side of both Tpm1.6 (Fig. 2B) and Tpm3.2 (Fig. 2C) is negatively charged. A notable exception is the N-terminus, which is positively charged in both tropomyosin isoforms (Fig. S1).

In the energy minimized Rosetta models, the actin residues Lys326, Lys328, and Asp311 were observed to interact with both Tpms in most of the actin-binding sites (in total 7 for Tpm1.6 and 6 for Tpm 3.2; Fig. 2; Table 1). Lys326 and Lys328 interact with side chains of two glutamate residues (in one instance the side chain of an aspartate residue) in one of the two Tpm α-helices, separated approximately by one turn of the α-helix (four residues apart). The models also revealed another type of salt bridge, where Asp311 on actin interacts with a lysine residue on the Tpm.

**Table 1.** Predicted salt bridges between actin and tropomyosin (Tpm) isoforms.

| Position | Actin | Tpm1.6 | Chain | Tpm3.2 | Chain |
| --- | --- | --- | --- | --- | --- |
|  | Residue | Residue |  | Residue |  |
| 1 | D311 | K248 | B | K212 | B |
| 1 | K328 | E257 | A | E221 | A |
| 1 | K326 | - | - | - | - |
| 2 | D311 | - | - | - | - |
| 2 | K328 | - | - | - | - |
| 2 | K326 | - | - | - | - |
| 3 | D311 | - | - | K132 | B |
| 3 | K328 | E173 | A | E137 | A |
| 3 | K326 | E177 | A | E141 | A |
| 4 | D311 | - | - | - | - |
| 4 | K328 | - | - | E102 | B |
| 4 | K326 | E131+E138 | B | E106 | B |
| 5 | D311 | - | - | - | - |
| 5 | K328 | E96 | A | E60 | A |
| 5 | K326 | D100 | A | - | - |
| 6 | D311 | - | - | R12+K13 | A |
| 6 | K328 | E54 | B | - | - |
| 6 | K326 | D58 | B | D22 | B |
| 7 | D311 | K7 | B | N/A | N/A |
| 7 | K328 | - | - | N/A | N/A |
| 7 | K326 | - | - | N/A | N/A |

Are there alternative binding modes between actin and Tpm to those described above? We tested this by sliding the Tpm coiled-coil models through the Tpm cryo-EM density at steps corresponding to one α-helical turn (∼5.5 Å) and keeping the Tpm coil-coil interface fixed (see Methods). We calculated relative Rosetta binding energy for the actin–Tpm interface at each position (Fig. S5). The models described above (no translation) had the lowest interface energies, supporting the notion that Tpm prefers these binding modes. After the Tpm chain had been translated by the distance of two actins (corresponding to 22 and 23 α-helical turns in Tpm1.6 and Tpm3.2, respectively), the same binding modes as described above were repeated. Sliding the Tpm sequences by the distance of one actin (corresponding to 12 α-helical turns in both Tpms), in turn, resulted in either the same (for Tpm1.6) or slightly higher (for Tpm3.2) energy. This alternate binding mode is feasible because the two α-helices in the coiled-coils alternate in making contacts to actin; in every second position, the other α-helix from the coiled-coil is proximal to actin. However, the phase of the α-helices differs between consecutive actin-binding sites (Fig. 2B,C). While in some positions the salt bridges are maintained, in other positions none or fewer of the actin sidechain rotamers are able to accommodate the shift in the Tpm α-helix, disrupting the salt bridges. The changing phase of the α-helix is also consistent with the absence of backbone signal in the cryo-EM density maps.

To conclude, our modeling not only predicts the same actin residues (Lys326, Lys328, and Asp311) that have been indicated in Tpm binding earlier (Lehman et al., 2019; Pavadai et al., 2020) but provides further insights into their role in binding. Lys326 and Lys328 exhibit at least two different rotamers, one pointing up and one down. These, together with the Tpm sequence at different actin interfaces, may provide enough flexibility to facilitate Tpm binding in different positions. Different Tpm sequences, vertical shift, and different sidechain rotamers likely accommodate the 5-Å lateral shift observed between the cryo-EM density between Tpm1.6 and Tpm3.2 tracks.

### A structure-based rationale for the absence of isoform mixing between Tpm1.6 and Tpm3.2

Earlier studies provided evidence that Tpm1.6 and Tpm3.2 cannot co-polymerize on the same actin filament, and thus segregate to different actin filament structures in cells (Gateva et al., 2017; Tojkander et al., 2011). The molecular basis behind these observations, however, has remained unclear. We addressed this question by using the actin:Tpm1.6 and actin:Tpm3.2 models. Placement of two Tpm1.6 coiled-coil Rosetta minimum-energy models consecutively on the same side of the actin filament allows the juxtaposition of the N-terminus from one dimer with the C-terminus of the next dimer (Fig. 3A). While the head-to-tail interaction is not revealed in our cryo-EM structures or Rosetta modeling, the juxtaposition shows that the phase of the Tpm1.6 α-helices and their positions match at the interaction site. The same is true for a homotypic juxtaposition of Tpm 3.2 coiled-coil with another Tpm 3.2 (Fig. 3B). In contrast, a heterotypic juxtaposition of Tpm1.6 and Tpm3.2 consecutively on an actin filament shows that neither the phase of the α-helices, nor their position matches those of the α-helices in the next coiled-coil (Fig. 3C,D). It is conceivable that both the phase of the α-helix and the position of the Tpm ends may play a role in promoting the formation of favorable head-to-tail interactions and further co-operative binding of the same isoform type on an actin filament. Thus, in addition to sequence-specific interactions between the N- and C-termini of Tpm dimers (Greenfield et al., 2006; Li et al., 2014; Rao et al., 2012), these geometrical factors indicated by our modeling are likely to play a crucial role in preventing heterotypic Tpm head-to-tail interactions. This hypothesis, based on the geometrical considerations and the observed 5-Å shift between Tpm1.6 and Tpm3.2 densities, requires further validation by cryo-EM of actin:Tpm1.6 and actin:Tpm3.2 complexes where the Tpm ends can be resolved. This could be possible by extending the approaches applied recently to visualize the muscle Tpm ends at low resolution in actin:Tpm:troponin complex (Yamada et al., 2020).

**Figure 3.**
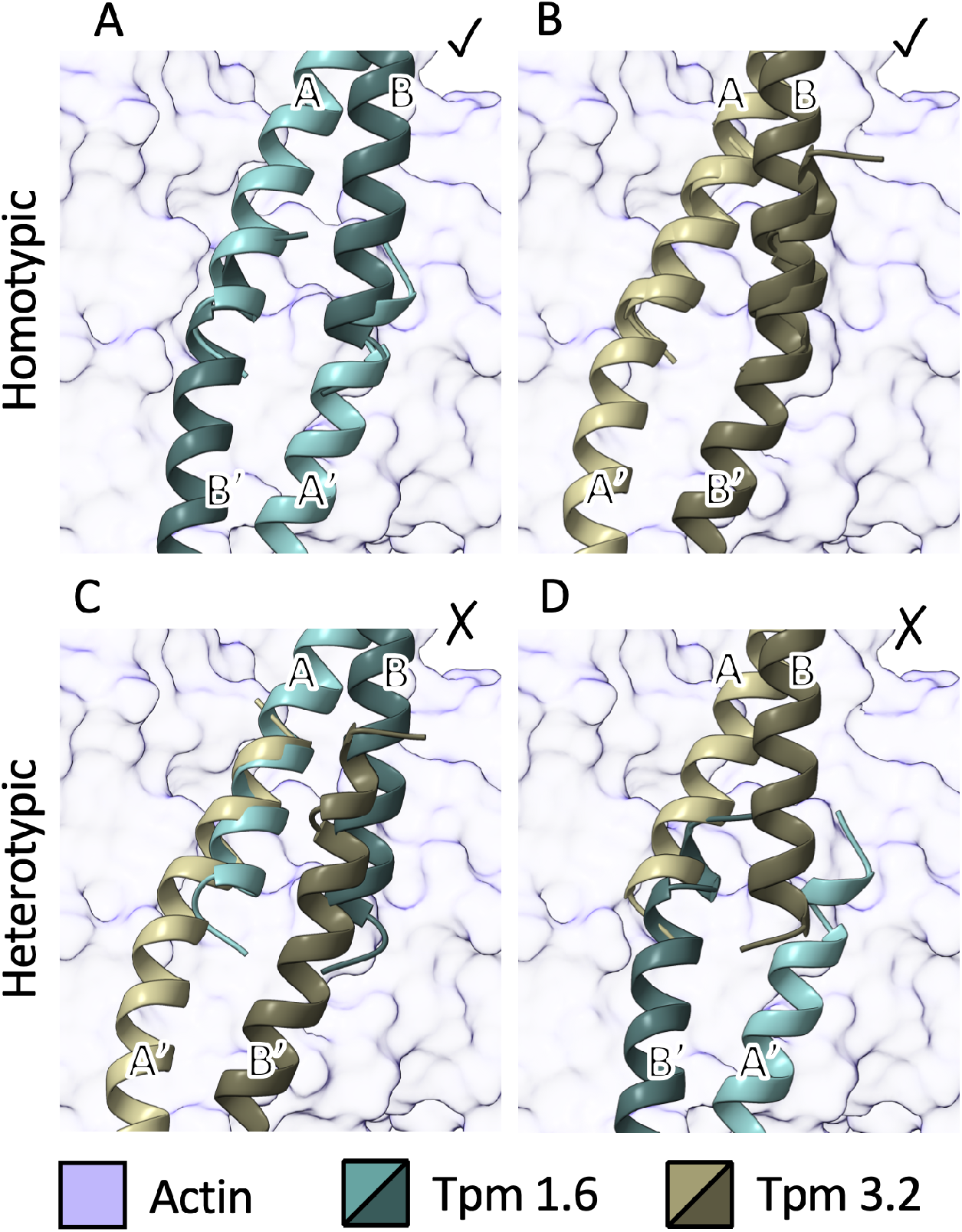
Visualisations of the tropomyosin ends. (**A, B**) Placement of two consecutive tropomyosins of the same kind (homotypic placement) are shown for Tpm1.6 (panel A) and for Tpm3.2 (panel B). (**C, D**) Placement of two consecutive tropomyosins of the different kinds (heterotypic placement) are shown. In all panels the two α-helices of the tropomyosin coiled-coil are labelled with A and B. The helices in the adjacent coiled-coil are labelled with A’ and B’. In all cases a conformational change in the Tpm ends would be required to allow binding of the next coiled-coil in the filament to avoid overlap. However, in the heterotypic complexes neither the phase of the α-helices, nor their position matches those of the α-helices in the adjacent coiled-coil.

### Comparison of structures reveals similarities and differences between non-muscle and muscle tropomyosin binding modes

We next compared the structures of actin:Tpm1.6 and actin:Tpm3.2 to the previously reported complexes between actin and muscle Tpms. The long non-muscle isoform Tpm1.6 occupies nearly the same position on actin as the skeletal muscle Tpm1.1 (Fig. 4A,B) (Von der Ecken et al., 2015), showing a shift of 2 Å in the direction of the filament long axis and rotation of 0.7 degrees around the axis. This ‘A-state’ position corresponds to the state of muscle Tpm on actin filament when no other proteins are present. In this context, it is important to note that muscle Tpm and non-muscle Tpm1.6 are encoded by the same *TPM1* gene, the only difference being that, as a result of alternative splicing, the last 27 amino acid residues of Tpm1.6 are different from those of Tpm1.1 (Fig. S1). In conclusion, this comparison shows that the C-terminal part of Tpm1 is not required for binding to the A-state position, but rather that both Tpm1.1 and Tpm1.6 bind the same position presumably via equivalent interactions. The C-terminal part, which is quite different between Tpm1.1 and Tpm1.6, may however play a role in preventing the mixing of different isoforms and/or promoting the binding of the next tropomyosin dimer via head-to-tail interactions.

**Figure 4.**
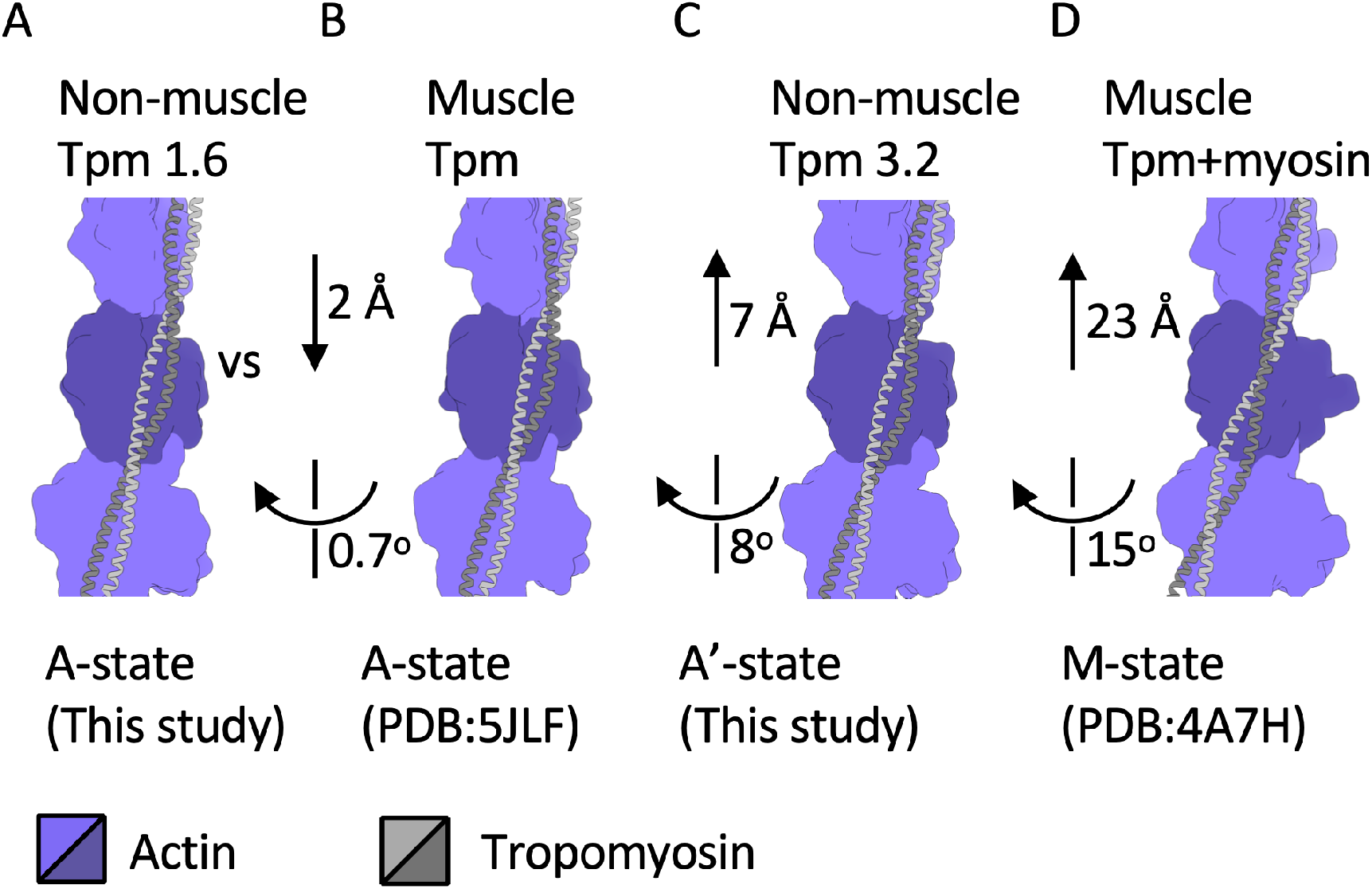
Comparison of the tropomyosin positions on the actin filament. (**A**–**D**) The position of Tpm1.6 in *A* (A-state) is compared to muscle tropomyosin in *B* (A-state), Tpm3.2 in *C* (A’-state), and to the one muscle tropomyosin from myosin bound (not shown) structure in *D* (M-state). The shifts and rotations shown indicate the measured transformations taking the tropomyosin coiled-coils in *B*–*D* on the reference coiled-coil in *A*.

The position of non-muscle Tpm3.2, in turn, is between the A-state and the myosin-bound M-state observed in the muscle actin:Tpm complex (Behrmann et al., 2012; Risi et al., 2021) (Fig. 4C,D). This state, denoted here as the A’-state, shows the Tpm3.2 shifted 7 Å along the filament long axis and rotated 8 degrees around it (the rotation corresponding to a horizontal shift of ∼6 Å; see Fig. 1E) relative to the muscle tropomyosin in the A-state (Figure 4C). For comparison, muscle Tpm in the M-state is shifted 23 Å and rotated 15 degrees relative to the A-state (Fig. 4D).

### Interplay of non-muscle tropomyosins with myosin II and ADF/cofilin

To evaluate if the different positions of Tpm1.6 (A-state) and Tpm3.2 (A’-state) on an actin filament affect their interactions with myosin II and ADF/cofilin, we carried out further binding experiments. An actin filament co-sedimentation assay demonstrated that ADP-loaded muscle myosin II S1 fragment binds with indistinguishable affinity to both Tmp1.6 and Tpm3.2-decorated actin filaments (Fig. S6A). Nevertheless, and similar to earlier experiments on non-muscle myosin II (Gateva et al., 2017), Tpm3.2-decorated actin filaments enhanced the actin-induced muscle myosin II ATPase activity when compared to bare actin or actin filaments decorated with Tpm1.6 (Fig. S6B). In conclusion, both non-muscle Tpm isoforms studied here allow myosin II binding to actin filaments, but only Tpm3.2 activates myosin II.

ADF/cofilin binds actin filaments in a cooperative manner and severs filaments at the boundaries between free and ADF/cofilin-decorated segments (Andrianantoandro and Pollard, 2006; Suarez et al., 2011; Wioland et al., 2017). ADF/cofilin and Tpms compete for actin binding, and Tpm isoforms protect actin filaments from ADF/cofilin-mediated severing to different extent (Gateva et al., 2017; Jansen and Goode, 2019; Ono and Ono, 2002). Interestingly, when the most abundant mammalian ADF/cofilin isoform, cofilin-1, is docked to Tpm–actin filaments by using earlier structural information (Tanaka et al., 2018), more pronounced steric clashes occur with Tpm1.6 than with Tpm3.2 (Fig. 5A-C).

**Figure 5.**
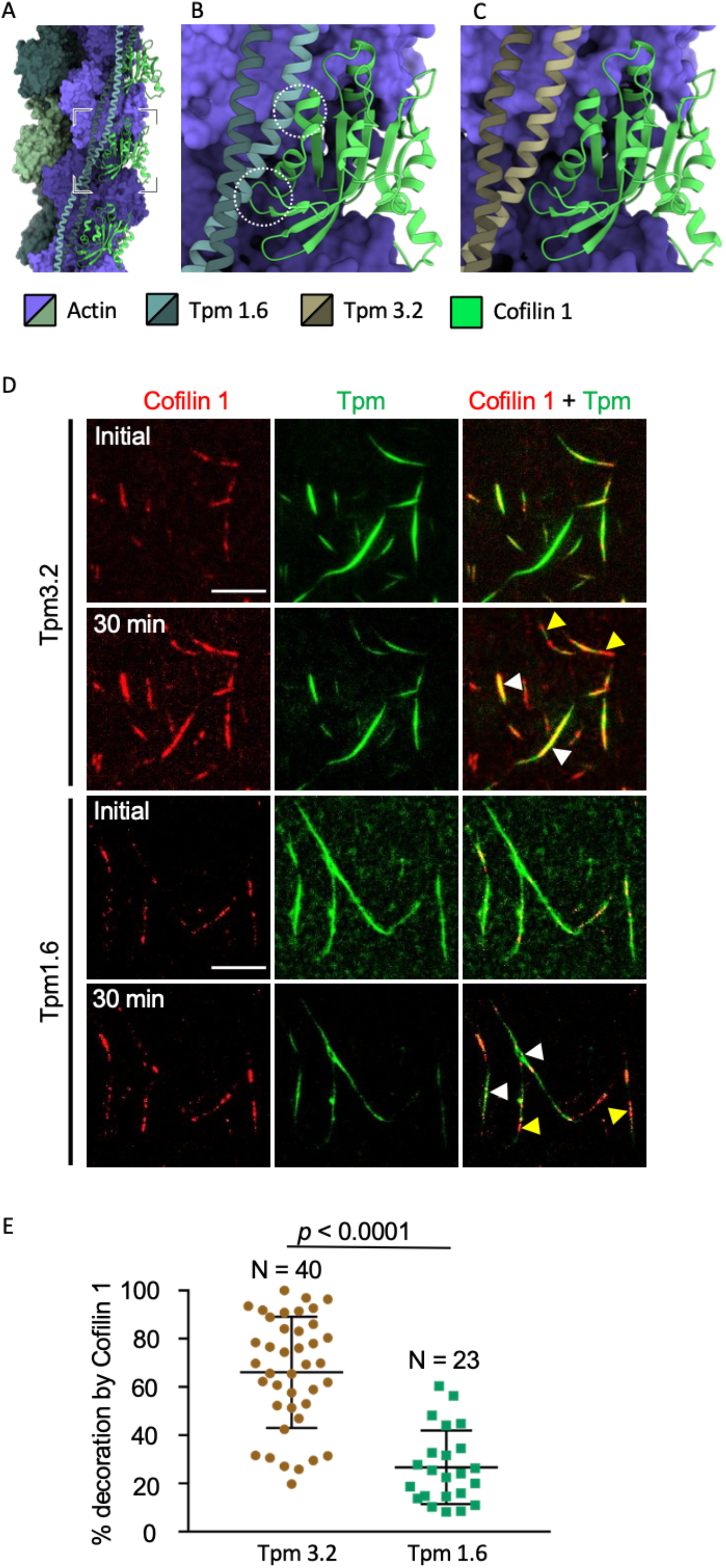
Influence of Tpm1.6. and Tpm3.2 on cofilin-binding to actin filaments. (**A**) A model of actin/Tpm1.6 complex with three copies of cofilin-1 (PDB:5YU8). (**B**) A close-up of the area indicated in *A*. Tpm1.6 binding site overlaps (dashed circles) with the one of cofilin-1. (**C**) The same view as in *B* is shown for actin/Tpm3.2 complex together with cofilin-1. No major clashes are evident. (**D**) Representative examples of *in vitro* TIRF microscopy showing mCherry-Cofilin-1 binding onto Tpm1.6 and Tpm3.2-decorated actin filaments. 1.2 μM sfGFP-TPm1.6 and 2.4 μM sfGFP-Tpm3.2 were mixed with pre-polymerized 0.8 μM actin and imaging was preformed after addition of 1.2 μM mCherry-Cofilin-1 (initial time point) and after 30 min incubation. White and yellow arrowheads indicate individual actin filaments and actin filament bundles, respectively. Tpm1.6-decorated actin filaments show less binding of cofilin-1 as compared to Tpm3.2 decorated filaments at 30 min time point. (**E**) The scatter plots show the proportion of cofilin-1 decoration on actin filaments in the presence of Tpm3.2 and Tpm1.6 at 30 min timepoint. Middle horizontal line = mean ± S.D from N=40 (Tpm3.2 from three technical repeats) and N=23 (Tpm1.6 from four technical repeats). The statistical significance between the groups was determined by using unpaired t-test with Welch’s correction.

To directly test ADF/cofilin-binding on actin filaments decorated by either with Tpm1.6 or Tpm3.2, we performed *in vitro* TIRF microscopy on sfGFP-fusions of Tpm1.6 and Tpm3.2, and mCherry-fusion of human cofilin-1. We first prepared sfGFP-Tpm-decorated actin filaments, mixed these with mCherry-cofilin-1, and imaged the samples immediately after mixing by TIRF microscopy. At our experimental conditions, we observed both individual actin filaments and filament bundles (Fig. 5D). Importantly, already at early time points, cofilin-1 segments were more visible on Tpm3.2-decorated actin filaments and filament bundles compared to Tpm.1.6-decorated filaments and bundles, and these differences became more pronounced after 30 min incubation (Fig. 5D,E). Thus, Tpm1.6 and Tpm3.2 inhibit actin filament binding of ADF/cofilin to different extents, and this may be at least partially due to the different actin filament binding interfaces of the two Tpm isoforms.

## DISCUSSION

Different non-muscle Tpm isoforms have distinct cellular functions, and they provide specific functional features to the actin filaments (Gunning et al., 2015a). However, the underlying structural principles have remained unknown. Moreover, the molecular basis by which different Tpm isoforms segregate to different actin filaments *in vitro* (Gateva et al., 2017) has remained elusive. Cryo-EM and biochemical experiments on non-muscle β/γ-actin filaments decorated by two non-muscle Tpm isoforms allowed us to probe the mechanisms underlying their specific functions. Most importantly, we revealed that while Tpm1.6 and Tpm3.2 binding leaves the actin conformation unaltered, they follow different paths along the major groove of the actin filament. We speculate that this property of Tpm contributes to their different biochemical functions and specific role in cells.

The different positioning of Tpm1.6 and Tpm3.2 along the actin filament provides a plausible explanation for why these two Tpm isoforms cannot co-polymerize with each other on actin filaments. This is because the heterotypic juxtaposition of Tpm1.6 and Tpm3.2 on an actin filament shows that neither the phase of the α-helices, nor their positions match with those of the α-helices in the adjacent tropomyosin coiled-coil. Therefore, whereas the N- and C-termini of Tpm dimers of the same isoform can generate proper head-to-tail oligomers, the N- and C-termini of Tpm1.6 and Tpm3.2 isoforms are out of the registry for head-to-tail interactions. In addition to these geometrical factors, also sequence-specific interactions between Tpm N- and C-termini contribute to their co-polymerization on actin filaments. However, because the C-terminal regions of Tpm1.6 and Tpm3.2 are nearly identical to each other, and for example much more similar than the ones between Tpm1.1 and Tpm1.6 (Fig. S1), we speculate that in the case of Tpm1.6 and Tpm3.2 the lack of isoform mixing must result mainly from their different positioning along the actin filament. In the future, it will be interesting to examine if also other most abundant non-muscle Tpm isoforms are ‘out of register’ with each other on an actin filament, as well as to study the specific roles of N- and C-terminal sequences of Tpms in their co-polymerization.

Earlier biochemical studies provided evidence that Tpm1.6 displays a much more stable association with actin filaments *in vitro* compared to Tpm3.2 (Gateva et al., 2017). This may be partially due to the presence of seven actin-binding sites in Tpm1.6 vs. six such sites in Tpm3.2. Additionally, Tpm1.6 occupies the ‘A-state’ position on an actin filament, whereas Tpm3.2 is shifted towards the M-state position, which at least in the case of muscle Tpm is energetically unfavorable. Because the central regions of Tpm1.6 and Tpm3.2 are highly conserved between each other and with muscle Tpm1.1 (Fig. S1), it is likely that the variable regions adjacent to N- and C-termini are responsible for the different positioning of Tpm1.6 and Tpm3.2 on the actin filament. Moreover, possible differences in the head-to-tail interactions may be responsible for the slow off-rate of Tpm1.6 from actin filaments as compared to the relatively dynamic association of Tpm3.2 with actin filaments. It is also important to note that our work, as well as earlier biochemical studies, were performed using Tpms produced in *E*.*coli*. These Tpms contain an acetylation-mimic Met-Ala-Ser sequence in the N-terminus. Tpms purified from mammalian cells that harbor native post-translational modifications, however, display slightly different affinities on actin filaments (Carman et al., 2021). Thus, also the dynamics of native Tpms on actin filaments may exhibit small differences from the ones produced in *E. coli*.

Our study sheds light on the different effects of Tpm1.6 and Tpm3.2 isoforms on ADF/cofilin and myosin II. We propose that, due to the larger overlap between Tpm1.6 and ADF/cofilin on actin filament, as compared to Tpm3.2 and ADF/cofilin, Tpm1.6 prevents more efficiently ADF/cofilin-binding to actin filament (Fig. 5). This could explain why Tpm1.6 protects actin filaments more efficiently from ADF/cofilin-mediated disassembly as compared to Tpm3.2 (Gateva et al., 2017). On the other hand, myosin II motor domain appears to bind both Tpm1.6-decorated and Tpm3.2-decorated actin filaments with similar affinity (Fig. S5), despite the fact that there seems to be a larger overlap between myosin and Tpm1.6 than between myosin and Tpm3.2 on an actin filament (Fig. 5). Thus, we propose that the differences in actin-mediated activation of myosin II ATPase activity between Tpm1.6 and Tpm3.2-decorated actin filaments are not due to their different binding sites along actin filaments, but rather result from sequence-specific interactions between Tpm3.2 and myosin II motor domain that enhance ATPase activation of the myosin. This is also supported by earlier work demonstrating that myosin-binding can shift muscle Tpms to a different position on the actin filament (Behrmann et al., 2012; Risi et al., 2021; Von der Ecken et al., 2015).

In conclusion, our work suggests that both the positioning of the Tpm isoform on actin filament, as well as sequence-specific interactions between Tpms and other actin-binding proteins, contribute to specific functions of different Tpm isoforms. In this context, it is important to note that non-muscle Tpms have so far been studied only in the context of a handful of actin-binding proteins, but they are likely to also control the activities of a large array of other actin-binding proteins, including the many members of the myosin family. Thus, more work is required to understand how non-muscle Tpms regulate the interactions with various cytoskeletal proteins with actin filaments. Moreover, it will be important to determine the structures of other non-muscle actin– Tpm filaments and uncover the molecular basis of their distinct biological and biochemical activities. Finally, it will be interesting to determine the high-resolution structures of native non-muscle Tpms to uncover the precise mechanisms by which properly post-translationally modified Tpms form head-to-tail interactions on actin filaments.

## MATERIALS AND METHODS

### Protein purification

The plasmids for the structural and biochemical studies of tagged and non-tagged human tropomyosins Tpm1.6 and Tpm3.2 were available from the previous work (Gateva et al., 2017). Non-tagged Tpms were purified as per a published method (Janco et al., 2016). Briefly, non-tagged Tpm constructs were expressed in *E*.*coli* BL21-DE3 bacterial expression system in 1 L Luria Bertani broth (2X) containing ampicillin (100 μg/ml) and was grown until the O.D_600_ reached 0.7–0.8. The grown culture was further induced with 1 mM IPTG final concentration at 37°C for 3 h. Cells expressing untagged Tpms were harvested by centrifugation (4000× g, 4°C, 15 min, JS 4.2 rotor, J6-MI centrifuge, Beckman Coulter) and resuspended in lysis buffer (20 mM Na_2_PO_4_, 500 mM NaCl, 5 mM MgCl_2_, 1 mM DTT and 1× protease inhibitor cocktail). The cells were lysed thoroughly by sonicating on ice (4×30s) until the bacterial suspension turned comparatively transparent. The cell lysate was heated in a water bath set at 80°C for 8 min. The heat-treated lysate was cooled to room temperature in a water bath. The supernatant containing Tpms was cleared from denatured proteins and cell debris by centrifugation at 47,810× g, 4°C for 45 min (F21S-8×50 rotor, Sorvall LYNX 4000, Thermo Fisher Scientific). Tpms were acid precipitated from the supernatant at pH 4.7 using 2 M HCl. The precipitated materials containing Tpms was centrifuged at 3900× g, 4°C for 12 min (F21S-8×50 rotor, Sorvall LYNX 4000, Thermo Fisher Scientific). The pellet was resuspended in a resuspension buffer (100 mM Tris-Cl pH 7.5, 500 mM NaCl, 5 mM MgCl2, 1 mM DTT, 1 mM NaN3). To dissolve the pellet completely, the buffer was readjusted to pH 7.0 using 1 M NaOH. The acid precipitation was repeated two more times to remove impurities and to obtain a white pellet. The pellet was resuspended in the resuspension buffer and filtered using a 0.22-μm membrane filter. Dialysis was performed overnight in a dialysis buffer (20 mM Tris-HCl pH 7.5, 500 mM NaCl, 0.5 mM DTT) using a 10-kDa molecular cutoff membrane (#88243, SnakeSkin™ Dialysis Tubing, 10MWCO, Thermo Fisher Scientific). The next day, a 4-ml SP QFF anion exchange column was equilibrated with 4 column volumes of anion exchange wash buffer (20 mM Tris-HCl pH 7.5, 10 mM NaCl, 0.5 mM DTT) and the supernatant was loaded onto the column at 2 ml/min loading speed. The column was washed with 4 column volumes of wash buffer. The column was further equilibrated with anion exchange elution buffer (20 mM Tris, pH 7.5, 1 M NaCl, 0.5 mM DTT) and Tpm was collected in the flow through. Samples from peak fractions were collected and analyzed by SDS-PAGE based on which the fractions were pooled together. The pooled fraction was diluted 4 times with hydroxyapatite wash buffer (10 mM Na_2_PO_4_ pH 7.0, 1M NaCl, 0.5 mM DTT) and loaded onto an equilibrated hydroxyapatite column with 0.5 ml/min loading speed. The column was washed with 4 column volumes of hydroxyapatite wash buffer and the protein was eluted in a 10 mM to 240 mM phosphate gradient against hydroxyapatite elution buffer (240 mM Na_2_PO_4_ pH 7.0, 1 M NaCl, 0.5 mM DTT). The eluted fractions were analyzed by SDS-PAGE and the fractions containing Tpm protein were pooled together. The acidification procedure was repeated twice to obtain pure protein. The precipitate was pelleted at 3900× g, 4°C for 12 min. The pellet was resuspended in dialysis buffer (20 mM Tris-HCl pH 7.0, 100 mM KCl, 5 mM MgCl_2_, 0.5 mM DTT, 0.02% NaN_3_) and dialyzed overnight at 4°C. The supernatant containing pure protein was concentrated using 30 MWCO column concentrator (Amicon® Ultra-15, Merck Millipore). Purified protein concentration was measured using spectrophotometer (NanoDrop® ND-1000 spectrophotometer, Thermo Fisher Scientific), fluorimetry and densitometry methods. The protein aliquots were flash frozen in liquid nitrogen and stored at –80°C.

The His-tagged sfGFP and mCherry fused tropomyosin and Cofilin-1 constructs were expressed in *E*.*coli* BL-21 (DE3) cells in 1 L auto-induction media at 37°C till O.D_600_ reached 0.7–0.8 after which the cells were induced at 28°C for 24 h. The cells were harvested by centrifugation at (4000× g, 4°C, 15 min, JS 4.2 rotor, J6-MI centrifuge, Beckman Coulter) and resuspended in Ni-NTA binding buffer (50 mM Tris-HCl pH 7.5, 300 mM NaCl, 10 mM imidazole, 1 mM DTT) and 1× protease inhibitor cocktail. Cells were lysed by sonication and cell debris was pelleted at 4000× g, 4°C for 15 min (JS 4.2 rotor, J6-MI centrifuge, Beckman Coulter). The his-tagged Tpm1.6 and Tpm3.2 were purified with Ni-NTA agarose beads (Qiagen) as per the manufacturer’s protocol. Ni-NTA agarose beads were first washed 2 times with Ni-binding buffer and centrifuged at 3000× g, 4°C for 5 min and the beads were suspended in equal volume of binding buffer. The Ni-NTA beads were incubated with the supernatant containing His-tagged fusion Tpm and mixed on an orbital shaker at 4°C, 1 h. The beads were loaded on a plastic column and washed extensively with low salt and high salt binding buffer and eluted in elution buffer (50 mM Tris-HCl pH 7.5, 300 mM NaCl, 250 mM imidazole, 1 mM DTT). The wash and eluted fractions were analyzed by SDS-PAGE. The elutes were dialyzed in dialysis buffer (10 mM HEPES pH 8.0, 20 mM NaCl, 1 mM DTT), overnight at 4°C. The supernatant was concentrated and was purified by gel filtration against gel filtration buffer with high salt (10 mM HEPES pH 8.0, 500 mM NaCl, 1 mM DTT). The peak fractions were analyzed by SDS-PAGE, pooled together and concentrated. The protein concentration was measured by fluorimetry, and the aliquots were flash frozen and stored at –80°C.

### Co-sedimentation assays

Actin co-sedimentation assays were performed as previously reported with few modifications (Doran et al., 2020). Non-muscle actin (β/γ-actin from human platelets) and myosin motor protein (S1 fragment) were procured from Cytosleleton Inc. and were resuspended in distilled water and stored at –80°C as per the manufacturer’s instructions. For competition assays, different amounts of either β/γ-actin or β/γ-actin and non-tagged Tpm1.6 or Tpm3.2 were mixed in presence of G-buffer (5 mM HEPES pH 7.4, 0.2 mM CaCl_2_, 0.2 mM DTT and 0.2 mM ATP). To completely saturate Tpm binding site on non-muscle actin filaments, actin and Tpm1.6 or Tpm3.2 were mixed in 4:1 ratio. Actin/ actin: Tpm were pre-polymerized by addition of F-buffer containing 20 mM HEPES pH 7.4, 100 mM KCl, 5 mM MgCl_2_, 0.2 mM EGTA, 1 mM DTT and 0.2 mM ATP final concentration at room temperature for 30 min. One μM of myosin motor protein was added to the polymerized actin/actin:Tpm complex and was incubated at room temperature for 30 min. Actin:Tpm:myosin motor protein complex was sedimented by centrifugation at 59000 rpm for 60 min at 10°C (TLA100 rotor, Beckman Optima MAX Ultracentrifuge). Supernatant and pellet fractions were separated, and samples were prepared for SDS-PAGE analysis by cooking in Laemmli Buffer at 100°C for 5 min. Supernatant and pellets were run on 4–20% gradient SDS-PAGE gels in equal proportions (Mini-PROTEAN TGX Precast Gels, Bio-Rad Laboratories Inc.). The gels were stained with Coomassie Blue staining solution (QC Colloidal Coomassie Stain, Biorad Laboratories, Inc.). The intensities of myosin motor protein bands was quantified with QuantityOne program (Bio-Rad), analyzed and plotted as amount of myosin motor protein in pellet (μM) to actin/ actin:Tpm1.6/ actin:Tpm3.2.

### Myosin ATPase assay

The steady-state ATPase activity of skeletal muscle myosin motor protein was measured using EnzChek® Phosphate Assay Kit (Molecular probes, Inc.) as per the manufacturer’s instructions. Before polymerization, β/γ-actin and Tpms were mixed in 4:1 ratio and preincubated in G-buffer (5 mM HEPES pH 7.4, 0.2 mM CaCl2, 0.2 mM DTT and 0.2 mM ATP). β/γ-actin:Tpm reactions were prepared with and without the addition of myosin motor protein. In the reaction containing myosin motor protein, β/γ-actin:Tpms were mixed with myosin motor protein in 10:1 ratio. The polymerization of β/γ-actin and β/γ-actin:Tpms was initiated using F-buffer (20 mM HEPES pH 7.4, 100 mM KCl, 5 mM MgCl_2_, 0.2 mM EGTA, 1 mM DTT and 0.2 mM ATP final concentration) at room temperature for 30 min. ATPase assay reaction mix was prepared by adding distilled water, actin-polymerization buffer, 0.2 μM substrate 2-amino-6-mercapto-7-methylpurine riboside (MESG), 1 U purine nucleoside phosphorylase (PEP), 0.6 μM myosin motor protein and incubated for 10 min at room temperature. The mix was transferred to a 96-well microreader plate. The reaction was started by adding 6 μM actin/actin:Tpm, mixed properly and the measurement was started without delay at 340 nm in a microplate reader at room temperature. All combinations were tested with and without the addition of myosin motor protein, Tpm1.6 and Tpm3.2. The myosin ATPase rate was calculated by the change in absorbance in presence of bare actin filaments and actin filaments decorated with Tpm1.6 or Tpm3.2 and plotted using GraphPad Prism 7 software. The data was analyzed for significance by one-way ANOVA followed by Tukey’s HSD post-hoc analysis.

### *In-vitro* TIRF microscopy

The TIRF chamber was pre-treated with 1.5% BSA for 1 h in a humidified chamber. β/γ-actin, sfGFP-fused Tpm1.6, Tpm3.2 and mCherry-fused Cofilin-1 were diluted in G-buffer containing 5 mM HEPES pH 7.4, 0.2 mM CaCl_2_, 0.2 mM DTT and 0.2 mM ATP. For polymerization 0.8 μM β/γ-actin was mixed with 1.2 μM sfGFP-Tpm1.6 and 2.4 μM sfGFP-Tpm3.2 in F-buffer (10 mM HEPES pH 7.0, 50 mM Na-acetate, 3 mM MgCl_2_, 1 mM DTT and 0.2 mM ATP) and incubated at room temperature for 30 min. 1.2 μM of mCherry-cofilin-1 was added to the reaction, followed by 1% methylcellulose final concentration. The reaction was mixed and injected into the BSA pre-treated TIRF chamber. The imaging was started immediately without any delay. TIRF imaging was performed using ONI Nanoimager equipped with 100× Apo TIRF 1.49 NA oil objective, 1000-mW 561-nm and 1000-mW 640-nm lasers, sCmos camera and NimOS software. The time-lapse images were captured for 30 min with 10 s interval using 3% green and red laser power at 100 fps speed. The time lapse images were analyzed by Fiji image J software. Only single actin:Tpm filaments were measured during the analysis. The growth of cofilin-1 segments in 30 min was analyzed by measuring the length of cofilin-1 segments at the first and the last frame of imaging and subtracting the length obtained at first frame from the last one. Thus, the rate of cofilin-1 extension on Tpm1.6 and Tpm3.2 decorated actin filaments was obtained by dividing the cofilin-1 segments by imaging time (30 min).

### Cryo-EM sample preparation, data collection and image processing

To dissolve the oligomers that form during the storage, 50 μM β/γ-actin was mixed in G-buffer (5 mM HEPES pH 7.4, 0.2 mM CaCl_2_, 0.2 mM DTT, 0.2 mM ATP), and incubated at room temperature for at least 30 min. To ensure complete saturation of actin filaments with Tpm, a 1:1.5 ratio of β/γ-actin:Tpm1.6 and Tpm3.2 was used for the sample preparation. β/γ-actin (12.5 μM) was mixed to F-buffer (10 mM HEPES pH 7.0, 50 mM Na-acetate, 3 mM MgCl_2_, 1 mM DTT and 0.2 mM ATP) and incubated at room temperature for 30 min to allow formation of actin filaments. Either Tpm1.6 (9 μM) or Tpm3.2 (9 μM) was added, mixed gently, and incubated further for 1 h at room temperature. The actin:Tpm complex was pelleted by centrifugation at 59,000 rpm for 1 h at 10°C (TLA100 rotor, Beckman Optima MAX Ultracentrifuge). The supernatant was discarded and fresh F-buffer containing 0.02% NaN_3_ was added to the pellet before incubating it overnight for the pellet to solubilize.

A 3-μl aliquot of the actin-Tpm sample was applied onto a glow discharged holey carbon copper grid (Quantifoil R1.2/1.3) and allowed to settle for 15 seconds. The grids were prepared using a vitrification apparatus (Vitrobot, Thermo Fisher Scientific) operated at a relative humidity of 95% and at 6°C. Grids were blotted for 5 s with filter paper prior to plunging into liquid ethane. The grids were stored in liquid nitrogen for subsequent screening and imaging.

Data were acquired using a transmission electron microscope (Talos Arctica, Thermo Fisher Scientific) equipped with Falcon III direct electron detector (Thermo Fisher Scientific). Movies of 45 frames were collected with a total electron exposure of 45 e^−^/Å^2^, with a defocus ranging from 1 to 3 μm. A total of 1,758 movies were recorded for Tpm1.6 and 1,273 movies for Tpm3.2 complex. The movies were then imported to RELION for data processing. The movies were aligned and summed using MotionCorr (Zheng et al., 2017). CTF was estimated using CTFFIND4 (Rohou and Grigorieff, 2015). Filaments were manually picked to generate a template for automated picking using 2D classification in RELION (Zivanov et al., 2018). Class averages showing Tpm-decorated filaments were used as templates and filaments were picked automatically using a helical rise of 27.5 Å and a width of 100 Å. Overlapping filament segments were extracted in 400×400-pixel boxes and subjected to reference-free 2D classification. Segments corresponding to classes representing bare actin were discarded and the remaining segments were subjected to two rounds of 2D classification. This resulted in 118,164 segments for Tpm1.6 and 100,804 segments for Tpm3.2. An ab initio model was generated for both datasets in RELION. Using this model as a reference, the filament segments were subjected to 3D classification applying helical symmetry. Particles corresponding to 3D class averages displaying Tpm decoration were selected. This resulted in 79,298 segments for Tpm1.6 and 81,431 segments for Tpm3.2. The segments were subjected to 3D reconstruction, using one of the 3D class averages as a reference. Helical symmetry was used during refinement and was allowed to explore the range from –161 to –170 degrees of rotation and a rise from 22 to 30 Å. After particle polishing, CTF refinement, and postprocessing, the final resolution was 3.9 Å for Tpm1.6 and 4.5Å for Tpm3.2 complex, as estimated by Fourier shell correlation (FSC=0.143). For visualizing the entire length of the actin filament corresponding to Tpms, the map was averaged in real space in a 512×512-pixel box by applying helical symmetry in RELION.

### Atomic model building and molecular modelling

A comparative model was created for β-actin in SWISS-MODEL (Waterhouse et al., 2018) using a fiber actin structure (PDB:5JLF) as a template. The resulting structure was fitted into the actin:Tpm cryo-EM density maps as a rigid body using UCSF ChimeraX (Pettersen et al., 2021). The structure was manually adjusted in Coot (Casañal et al., 2020) and in ISOLDE (Croll, 2018). Symmetry copies (8 chains in total) were generated in ChimeraX by rigid body fitting. A poly-alanine model (135 residues, from PDB:3J8A) of Tpm was fitted into the density maps filtered to 7 Å resolution as a rigid body in ChimeraX and refined in ISOLDE in the same density maps using secondary structure restraints. The Tpm models were combined with their respective actin models and the combined models were refined in real space in Phenix (Afonine et al., 2018) against the unfiltered maps, using the parameters created in ISOLDE with command *isolde write phenixRsrInput* with added non-crystallographic symmetry constraints. Structures were validated using Phenix and MolProbity (Williams et al., 2018). Figures were made using ChimeraX.

Modeling of the actin:Tpm interfaces was carried out using a combination of AlphaFold structure prediction (Jumper et al., 2021) and Rosetta all-atom refinement (Song et al., 2013; Wang et al., 2016). For each actin filament, the model was first refined symmetrically into the corresponding cryo-EM density map. These refined actin filament models were then used in the subsequent modeling of Tpms. To identify the contacts between the Tpm and actin interface, the following procedure was performed on both Tpm1.6 and Tpm3.2:

1. Prediction of the Tpm coiled-coil by AlphaFold: A model of the coiled-coil alone was predicted by AlphaFold, using Tpm monomer multiple sequence alignments with a depth of over 2000 sequences. Five models from AlphaFold were minimized in Rosetta using the *Relax* protocol (Conway et al., 2014) and the lowest-scoring model was chosen. For both sequences, the final model had at least 85% residues with a confidence estimate (pLDDT) greater than 70. Predicted alignment error (PAE) was less than 5 Å for at least 60% of contacts made across the interface of the coiled-coil.
2. Identification of favorable actin– Tpm interfaces: We first identified favorable interactions of actin and Tpm by modeling the energetics of a small piece of Tpm (approximately 45 residues from each Tpm chain) interacting with a single actin monomer. We considered all sequence registrations onto this helix that maintained the predicted coiled-coil interface, by first shifting the helices “up” by one turn (equivalent to a 3.5-residue shift), and then by rethreading the model offset by 7 residues (equivalent to a shift of one heptad repeat). Each model was refined with Rosetta’s *Relax* function.
3. Determination of the full-length actin–Tpm interface: We translated the full-length Tpm model in relation to the actin filament in steps corresponding to one helical turn (approximately 5.5 Å) and refined the resulting model. Refinement involved a two-step procedure. Since the predicted Tpm models showed different curvature than was indicated by the density, we first refined Tpm in isolation (with Rosetta’s *Relax*) using fit-to-density as well as constraints from the interfaces determined in step 2. Next, we refined the full actin:Tpm complex, using the fit-to-density function without any other constraints (again using *Relax*). In this step, only actin residues within 8 Å of Tpm were permitted to move. For this latter step, five independent trajectories were run. For each position, the interface energies were calculated using the *ddG* application in Rosetta (Fig. S5).
4. Final refinement of full actin:Tpm complex: The model from step 3 with the lowest interface energy was refined using Rosetta’s *LocalRelax* protocol (Wang et al., 2016). Twenty models were generated, and the lowest energy model was selected, using Rosetta’s force field augmented with the “fit to density” score term.

## ACKNOWLEDGEMENTS

We thank Pasi Laurinmäki and Benita Löflund for technical assistance in cryo-EM. The facilities and expertise of the HiLIFE cryo-EM unit at the University of Helsinki, a member of Instruct-ERIC Centre Finland, FINStruct, and Biocenter Finland are gratefully acknowledged. The authors wish to acknowledge CSC – IT Center for Science, Finland, for computational resources. This study was supported by grants from Academy of Finland (302161 to P.L.) and Sigrid Jusélius Foundation (4708344 to P.L.). Work in the laboratory of J.T.H. was supported by Helsinki Institute of Life Science HiLIFE.

## AUTHOR CONTRIBUTIONS

Conceptualization, F.D., P.L. and J.T.H.; Formal Analysis, M.S., T.K. and J.T.H; Investigation, M.S., S.K., R.G., K.K., T.K., K.E.; Writing – Original Draft, S.K., P.L. and J.T.H.; Writing – Review & Editing, M.S., S.K., P.L. and J.T.H.; Visualization, J.T.H.; Funding Acquisition, P.L.; Resources, S.K., E.K. and K.K.; Supervision, F.D., P.L. and J.T.H.

## DATA AVAILABILITY

Cryo-EM maps and models generated during this study are available from Electron Microscopy Data Bank (EMDB) accession codes EMD-14957 and EMD-14958, and Protein Data Bank (PDB) accession codes 7ZTC and 7ZTD.

## SUPPLEMENTARY FIGURES

**Supplementary Figure S1.**
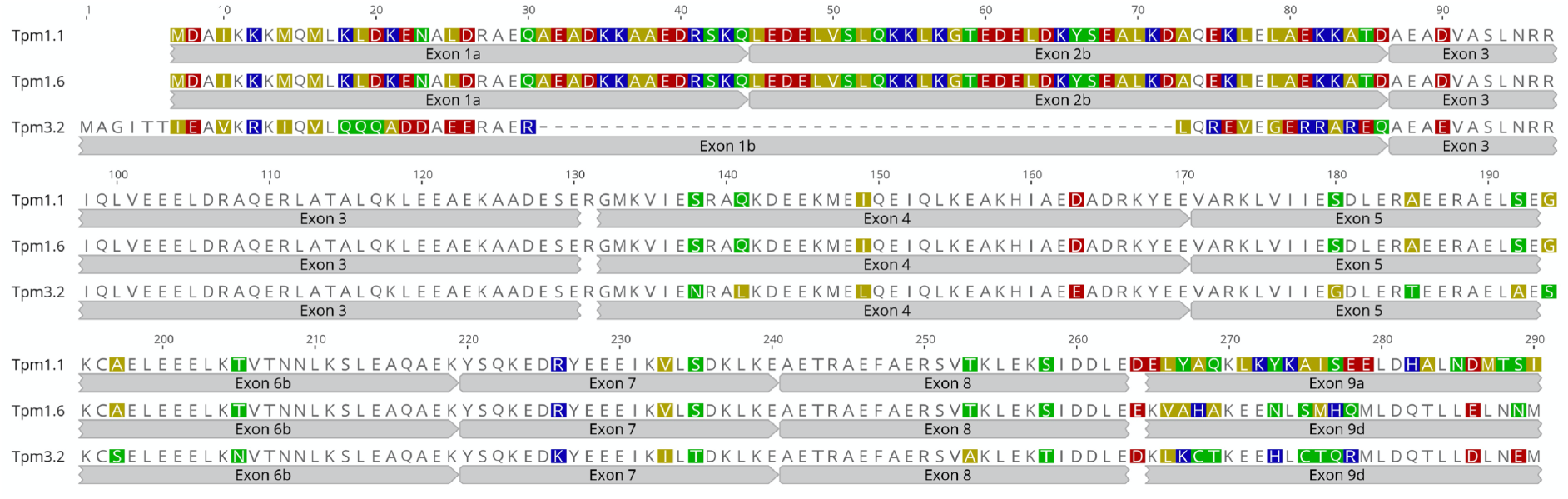
Amino acid sequences of muscle Tpm1.1 and non-muscle Tpm1.6 and Tmp3.2. The exon structures are shown below each sequence. The residues on the white background are conserved in all three tropomyosins, whereas the non-conserved positions are colored according to the charge of residue. The numbering corresponds to the alignment position. There is a gap of 43 residues in Tpm3.2 (results from the lack of exon 2) that roughly corresponds to a single actin molecule in the filament.

**Supplementary Figure S2.**
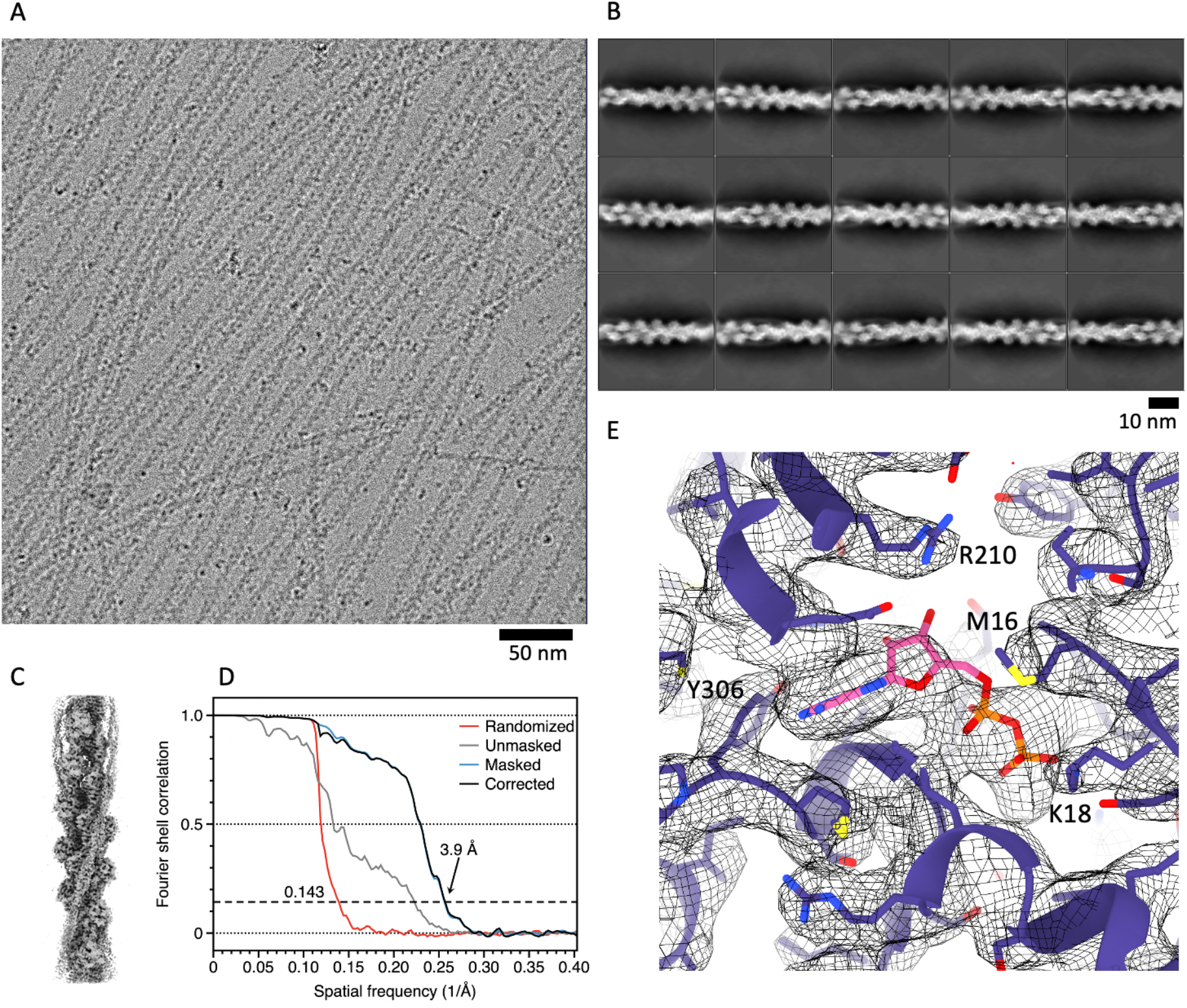
Cryo-EM analysis of the actin:Tpm1.6 complex. (**A**) A representative micrograph of actin:Tpm1.6 filaments. (**B**) Two-dimensional class averages showing Tpm1.6 decorated actin filaments. (**C**) Cryo-EM map of actin:Tpm1.6 complex. (**D**) Resolution assessment by Fourier shell correlation. The resolution, where the masking effect corrected curve (black), drops below the threshold (dashed line) is indicated with an arrow. (**E**) A close-up of the ATP-binding site in the actin:Tpm1.6 complex. Cryo-EM density is shown as a mesh. Some of the residues close to the ADP (pink) are labeled.

**Supplementary Figure S3.**
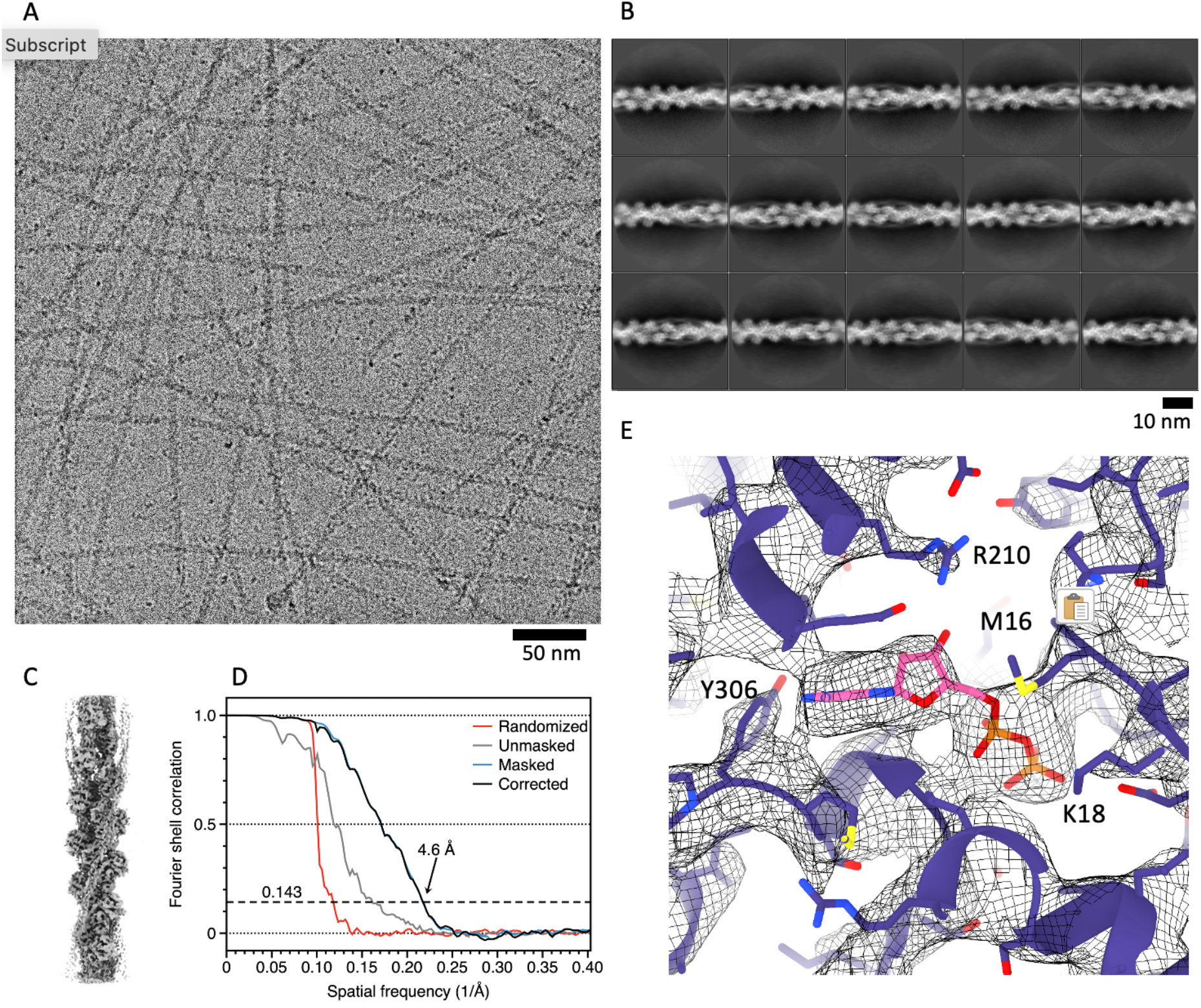
Cryo-EM of the actin:Tpm3.2 complex. (**A**) A representative micrograph of actin:Tpm3.2 filaments. (**B**) Two-dimensional class averages showing Tpm3.2 decorated actin filaments. Segments contributing to class-averages outlined in blue were used to refine the final map. (**C**) Cryo-EM map of actin:Tpm3.2 complex. (**D**) Resolution assessment by Fourier shell correlation. The resolution, where the masking effect corrected curve (black), drops below the threshold (dashed line) is indicated with an arrow. (**E**) A close-up of the ATP-binding site in the actin:Tpm3.2 complex. Cryo-EM density is shown as a mesh. Some of the residues close to the ADP (pink) are labelled.

**Supplementary Figure S4.**
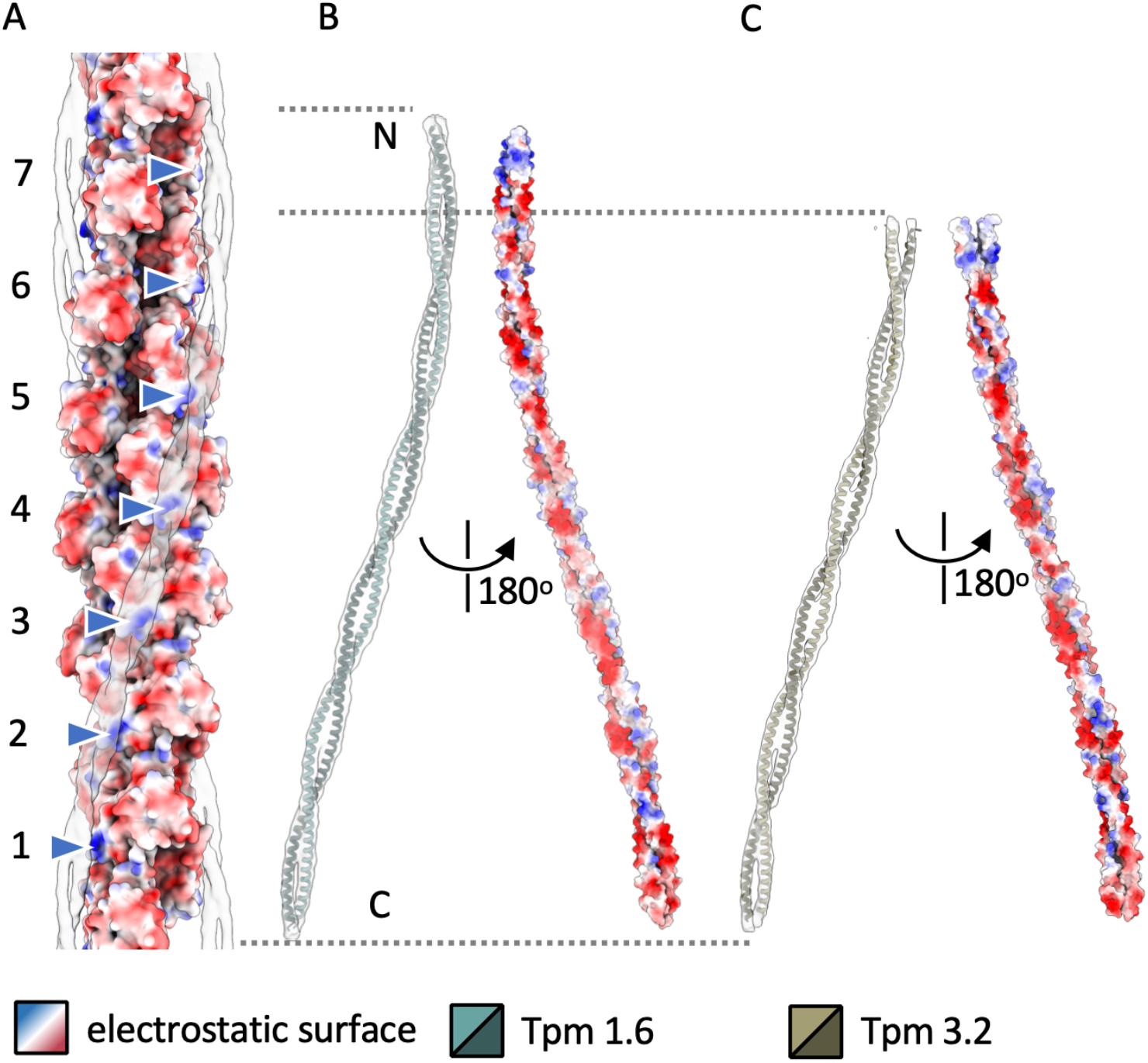
Electrostatic potential surface of actin and bound tropomyosins. (**A**) A surface rending or actin:Tpm1.6 complex is shown. The actin surface is colored based on its electrostatic potential (blue, positively charged; red, negatively charged). The position of a positively charged patch (blue arrowheads), consisting of residues K326 and K328 is shown in seven consecutive actin monomers (1–7). The cryo-EM density of Tpm1.6, filtered to 7 Å resolution, is shown as a transparent surface. (**B**) Model of Tpm1.6 is shown as a ribbon, overlayed on the cryo-EM density (transparent surface) on the left. The electrostatic surface of Tpm1.6 is shown rotated by 180 degrees on the right. The face opposing actin is largely negatively charged, the N-terminus (N) being an exception. (**C**) The same renderings as in *B* are shown for Tpm3.2.

**Supplementary Figure S5.**
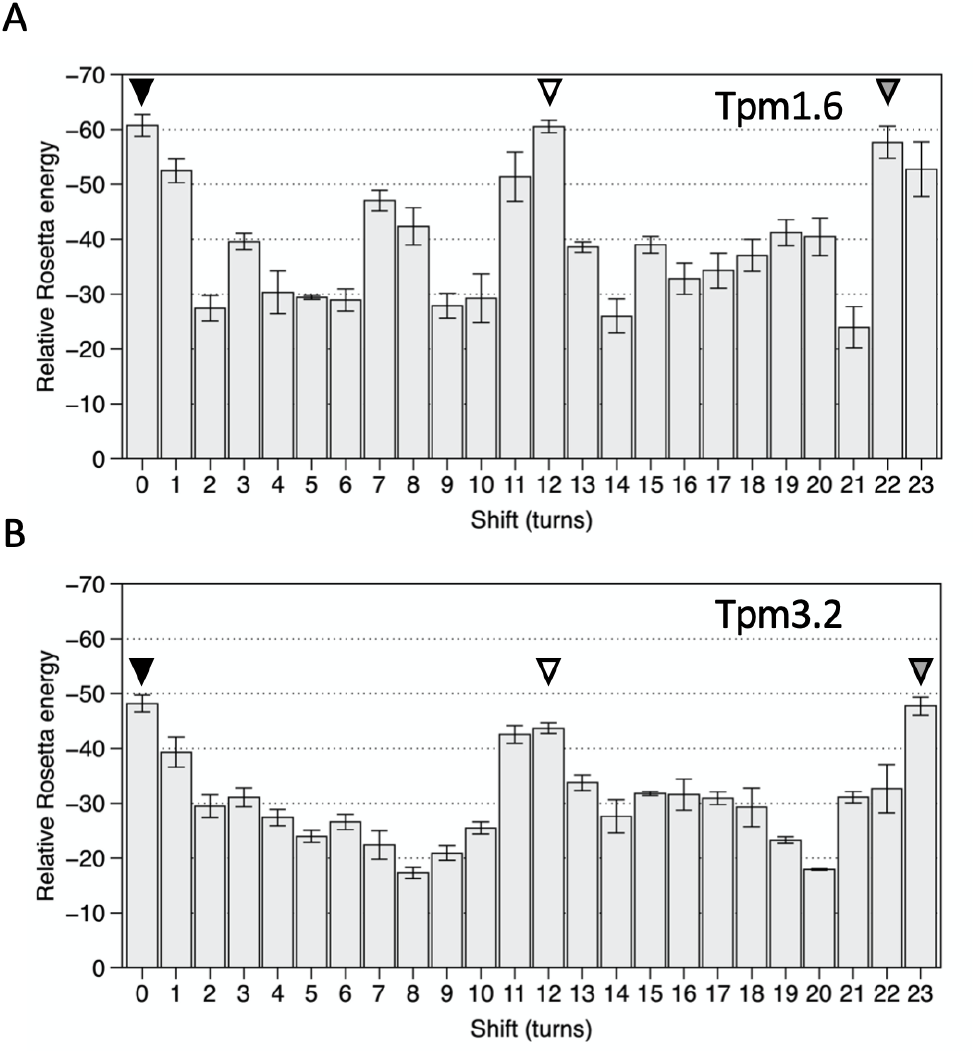
Rosetta energy of the tropomyosin models in different positions. (**A**–**B**) The relative Rosetta energy is plotted as a function of shift, determined as the number of alpha-helical turns, for Tpm1.6 in *A* and for Tpm3.2 in *B*. The error bars denote standard deviations calculated from five independent trajectories. The approximate positions of the energy minima are denoted at 0 turns (black triangle), at 12 turns (white triangle), and at 22 or 23 turns (gray triangle).

**Supplementary Figure S6.**
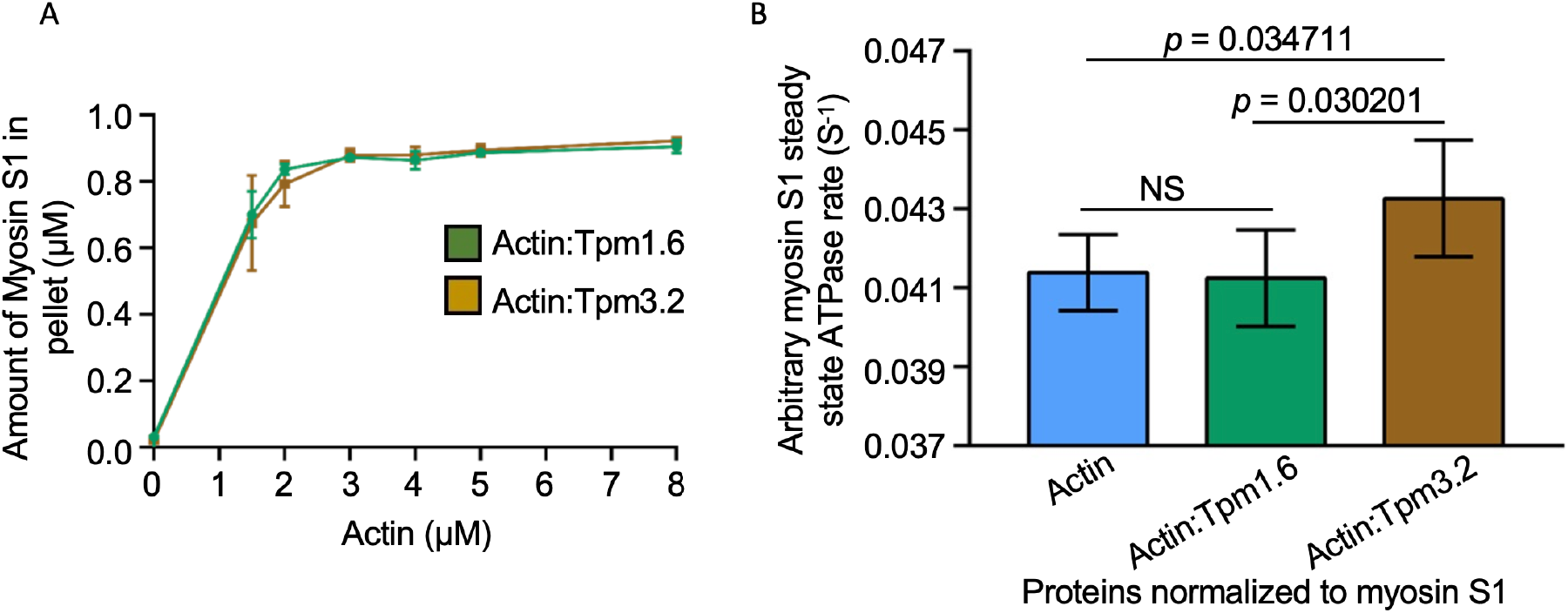
Effects of Tpm1.6 and Tpm3.2 on myosin II activation and actin binding. (**A**) Actin filament co-sedimentation assay demonstrating that skeletal muscle myosin II S1 fragment binds Tpm1.6- and Tpm3.2-decorated β/γ-actin filaments with similar affinity. Actin and Tpm1.6/Tpm3.2 were mixed in 4:1 ratio to ensure saturation of actin filaments by tropomyosins. The concentration of ADP-loaded myosin S1 fragment was 1 μM and actin concentration was varied from 0 to 8 μM. The line plot represents the median value from three independent experiments. Error bars ± S.D. (**B**) The skeletal muscle myosin S1 fragment steady state ATPase activity was measured in the presence of 6 μM bare β/γ-actin filaments or 6 μM Tpm1.6/Tpm3.2-decorated β/γ-actin filaments. The rate of ATP hydrolysis represents the arbitrary ATPase units (A.U.) per second. The myosin ATPase activity mean value was calculated from five independent repeats. The error bars represent S.E.M.

**Supplementary Table 1:** Cryo-EM data collection and refinement statistics.

|  | **Actin:Tpm1.6**<br>EMD-14957<br>PDB:7ZTC | **Actin:Tpm3.2**<br>EMD-14958<br>PDB:7ZTD |
| --- | --- | --- |
| **Data collection** |  |  |
| Magnification | 150,000 | 150,000 |
| Voltage (kV) | 200 | 200 |
| Defocus range ($\mu\text{m}$) | 1.0–3.0 | 1.0–3.0 |
| Pixel size ($\text{\AA}$) | 0.97 | 0.97 |
| Exposure rate ($\text{e}^-/\text{pix}/\text{sec}$) | 1 | 1 |
| Frames per exposure | 45 | 45 |
| Total electron exposure ($\text{e}^-/\text{\AA}^2$) | 45 | 45 |
| Micrographs (no.) | 1700 | 1250 |
| Helical symmetry imposed | rise: 27.8 $\text{\AA}$<br>turn: $-166.5^\circ$ | rise 27.8 $\text{\AA}$<br>turn $-166.4^\circ$ |
| Initial particle images (no.) | 118,164 | 100,804 |
| Final particle images (no.) | 79,298 | 81,431 |
| Map resolution ($\text{\AA}$) | 3.9 | 4.6 |
| FSC threshold | 0.143 | 0.143 |
| Map sharpening B factor ($\text{\AA}^2$) | $-147$ | $-170$ |
| **Refinement** |  |  |
| Initial model used (PDB code) | 5JLF, 3J8A | 5JLF, 3J8A |
| Model-to-map resolution ($\text{\AA}$) | 4.1 | 4.6 |
| FSC threshold | 0.5 | 0.5 |
| Model-to-map correlation | 0.82 | 0.80 |
| Model composition | 8 actin chains, 4 Tpm chains | 8 actin chains, 4 Tpm chains |
| Protein residues | Actin: residues 7–371<br>Tpm: 135 poly-Ala | Actin: residues 7–371<br>Tpm: 135 poly-Ala |
| Ligands | 1 ADP per actin | 1 ADP per actin |
| **R.m.s. deviations** |  |  |
| Bond lengths ($\text{\AA}$) | 0.004 | 0.007 |
| Bond angles ($^\circ$) | 0.843 | 1.042 |
| **Validation** |  |  |
| MolProbity score | 1.73 | 1.81 |
| Clash score | 5.29 | 8.02 |
| Rotamer outliers (%) | 0.00 | 0.00 |
| **Ramachandran plot** |  |  |
| Favoured (%) | 93.08 | 94.57 |
| Allowed (%) | 6.92 | 5.43 |
| Outliers (%) | 0.00 | 0.00 |

